# Targeted measurement of blood flow in the Anterior Cerebral Artery using Diffuse Correlation Spectroscopy

**DOI:** 10.64898/2026.07.25.740702

**Authors:** Susweta Das, Kavita Sharma, Soumyajit Sarkar, Kimberly Gonsalves, U.S. Srinivasan, Hari M Varma

## Abstract

**Significance:** Diffuse Correlation Spectroscopy (DCS) is an established technique for non-invasive monitoring of Cerebral Blood Flow (CBF), but existing applications primarily measure CBF changes averaged over cortical tissue volumes. A method capable of targeting blood flow within a particular intracranial artery would expand the utility of DCS for vessel-specific cerebral perfusion monitoring and continuous bedside assessment.

**Aim:** We aim to investigate the feasibility of targeted, non-invasive monitoring of blood flow in the Anterior Cerebral Artery (ACA) using DCS through optimization of probe geometry and placement.

**Approach:** A custom-built DCS system operating at 785 nm was used to probe ACA from the glabellar region. Source-detector (SD) separation, probe orientation, and probe location were systematically optimized using lower-limb motor tasks and a mental arithmetic task. The optimized configuration was evaluated using an ACA-mimicking multilayer phantom and validated in forty healthy volunteers during lower-limb activation tasks and postural changes.

**Results:** An SD separation of 17 mm, vertical probe orientation, and glabellar placement provided the highest sensitivity to ACA-related blood flow changes. Phantom experiments demonstrated sensitivity to flow changes in a vessel located 45 mm beneath the scalp and showed that the frontal sinus, cerebrospinal fluid, and the absence of cortical tissue beneath the glabella along the longitudinal fissure together provide the best optical window for probing deep ACA flow. Using the optimized probe configuration, significant increases in relative CBF were observed during standing leg marching (66.4 ± 38.6%) and supine leg crunches (39.4 ± 32.2%) (p < 0.01), with consistent responses during supine-to-stand postural transitions.

**Conclusion:** The proposed DCS approach enables targeted measurement of blood flow within the ACA territory. This technique provides a framework for continuous, non-invasive monitoring of ACA perfusion and has potential applications in cerebrovascular monitoring and stroke care.

## 1 Introduction

Cerebral Blood Flow (CBF) is a critical physiological parameter that reflects the delivery of oxygen and nutrients to brain tissue and is therefore an important biomarker of cerebral health and function. Alterations in CBF are associated with several neurological disorders, including ischemic stroke, traumatic brain injury, and cerebrovascular diseases. In acute ischemic stroke, timely assessment of regional blood flow is particularly important because the extent and progression of tissue damage depends strongly on cerebral perfusion. Clinical management during the first 6–48 hours after stroke onset requires monitoring cerebral hemodynamics to assess disease progression and determine the need for timely therapeutic intervention [1]. Consequently, there is a significant need for techniques capable of continuously monitoring CBF at the bedside during this critical period [2].

Computed Tomography (CT) and Magnetic Resonance Imaging (MRI) are routinely used for diagnosing stroke and visualizing affected brain regions. While these modalities provide anatomical and perfusion information with a high degree of accuracy, they reflect changes that are taking place within the brain at the time of scan which may change as disease progresses over a period of time. Hence, they are impractical for repeated measurements especially in acute cases where the patient is on life support and are therefore unable to provide real-time measurements over extended periods [3]. As a result, they cannot capture the temporal evolution of cerebral blood flow occurring over hours following stroke onset.

Transcranial Doppler (TCD) ultrasound is a widely used non-invasive technique for measuring blood flow velocity in major cerebral arteries. Owing to its portability and high temporal resolution, TCD has found applications in stroke monitoring and assessment of cerebrovascular function [4]. However, TCD measurements are highly operator dependent and require appropriate expertise to obtain reliable data. Furthermore, successful insonation depends on the availability of an adequate acoustic window through the skull, which may be absent in a significant fraction of subjects [5]. These limitations make continuous bedside monitoring challenging, particularly in acute clinical settings. TCD measurements are most commonly performed in the Middle Cerebral Artery (MCA) through the transtemporal acoustic window. In contrast, assessment of the Anterior Cerebral Artery (ACA) is considerably more difficult because of its anatomical location and the increased thickness of the frontal bone [6, 7]. Consequently, direct and continuous non-invasive monitoring of ACA blood flow from the frontal region remains challenging using conventional ultrasound-based approaches.

Diffuse Correlation Spectroscopy (DCS) has emerged as a promising optical technique for non-invasive measurement of cerebral blood flow. DCS utilizes temporal fluctuations of near-infrared light scattered by moving red blood cells to quantify blood flow dynamics in deep tissue [8, 9]. The technique offers several advantages, including portability, continuous monitoring capability, and suitability for bedside applications. Previous studies have successfully employed DCS to investigate cerebral hemodynamic responses during functional activation tasks such as walking [10], finger tapping [11], mental arithmetic [12, 13] and voluntary apnea [14] where the measurements have been obtained primarily from the left or right prefrontal areas or the temporal cortex. DCS has also been used to study cerebral autoregulation during postural changes and head-of-bed manipulations [15, 16]. However, in these applications, DCS measurements typically represent the volumetric blood flow changes in the microvasculature within the probed cortical region rather than a specific major cerebral artery.

In this study, we investigated the feasibility of directly measuring blood flow in ACA through the glabella region using DCS, as demonstrated briefly in one of our previous works [17]. ACA is one of the principal terminal branches of the internal carotid artery and supplies the medial frontal and medial parietal lobes, orbitofrontal cortex, cingulate cortex, corpus callosum, basal ganglia, anterior hypothalamus and anteriorinferior portion of the internal capsule, paracentral lobule which includes lower limb motor and sensory representation [18, 19]. Although ACA strokes account for only a small proportion of ischemic stroke cases, occlusion of this vessel can produce neurological deficits including contralateral lower-extremity weakness, gait disturbances, behavioral changes, and cognitive impairment [20, 21]. Therefore a method capable of directly monitoring ACA blood flow non-invasively would be valuable for stroke assessment and management, especially in situations where conventional ultrasound approaches are limited.

To identify the optimal set of probe parameters to maximise the sensitivity towards the blood flow changes in ACA we initially took measurements with multiple source-detector (SD) separations (15 mm, 17 mm, and 20 mm) and SD orientations from the glabellar region in healthy subjects. Next in order to identify the most suitable measurement location, we measured the blood flow changes during lower-limb motor tasks and cognitive activation tasks from the glabella and the left pre-frontal region (Fp1). Statistical comparisons of the hemodynamic responses obtained from different frontal locations reveal that measurements acquired at the glabella provide the strongest and most consistent ACA-related signals. Now having determined the set of optimal probe configuration and location we initially tested these settings in an ACA mimicking multi-layer phantom to evaluate whether sufficient optical sensitivity is achieved at a depth of 45 mm which is the approximate depth of the A3 segment of ACA from the scalp surface [22].

Finally, using the optimized measurement configuration consisting of a glabellar probe placement and an SD separation of 17 mm in vertical orientation along the midline, we evaluated ACA blood flow responses in a cohort of 40 healthy subjects performing two lower-limb activation tasks and postural changes including supine and standing positions. The results show consistent increase in rCBF from stand to supine and a decrease from supine to stand position. Additionally, consistent and statistically significant increases in blood flow during task execution with respect to resting periods was seen that support the feasibility of direct, non-invasive ACA monitoring using DCS.

## 2 Methods and Materials

### 2.1 DCS instrumentation

The in-house developed DCS system used in this study is shown in Fig. 1. The system employed a fiber-coupled continuous-wave laser operating at 785 nm (linewidth < 0.1 nm, Holmarc Opto-Mechatronics Ltd., India) as the illumination source. Light was delivered to the tissue through a multimode optical fiber with a core diameter of 400 *µ*m. The diffusely scattered photons were collected at an SD separation of 15 mm, 17 mm and 20 mm separately, using a single-mode optical fiber and detected using a single-photon avalanche diode (SPAD; SPCM-AQRH-14-FC, Excelitas Technologies, USA).

**Fig 1.**
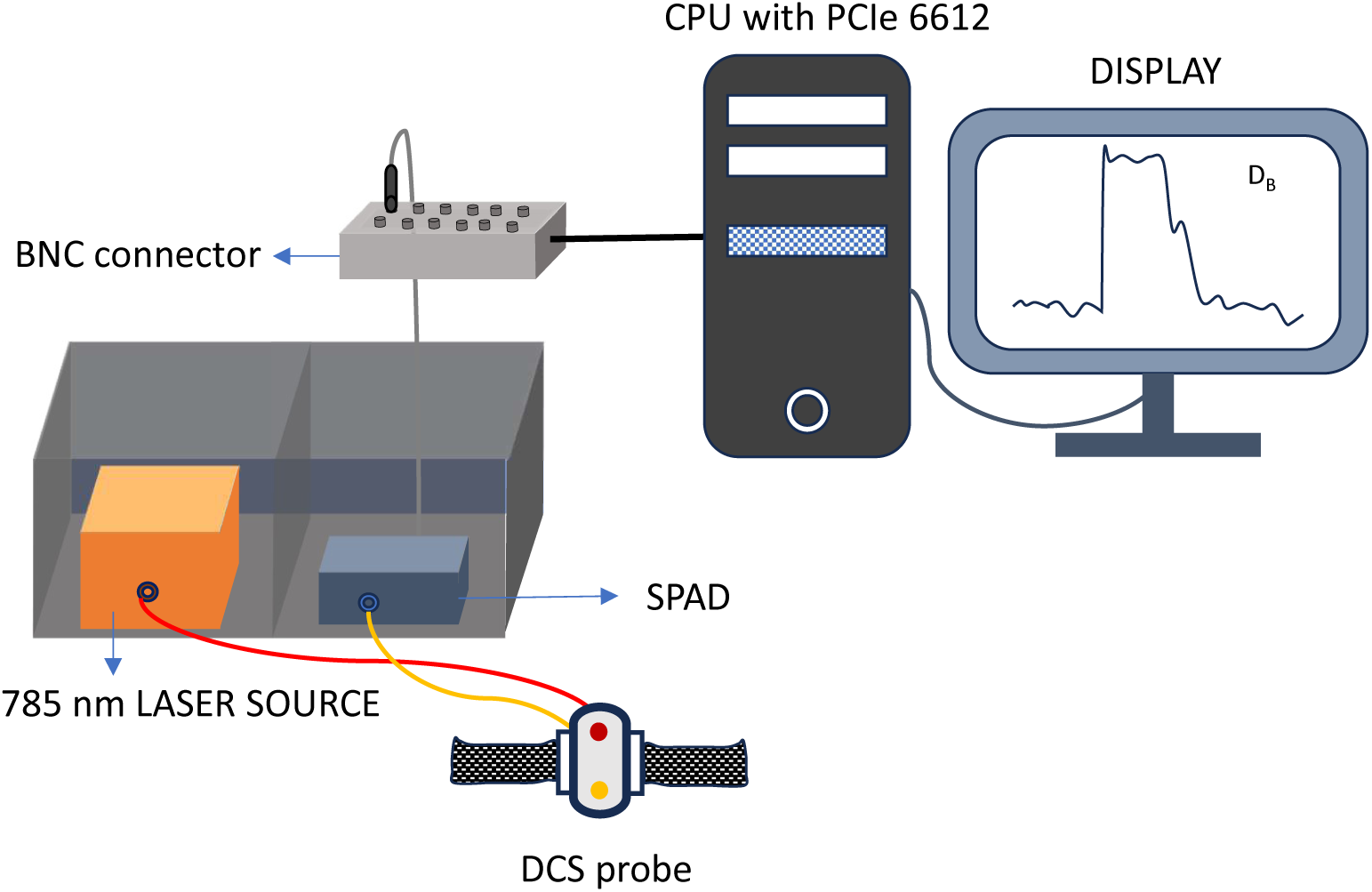
Schematic of the Diffuse Correlation Spectroscopy (DCS) system: A 785-nm fiber-coupled laser delivers light to the tissue through a multimode (MM) fiber integrated within the optical probe. The diffusely scattered light is collected by a single-mode (SM) fiber and detected using a single-photon avalanche diode (SPAD). The SPAD output is acquired by a data acquisition unit to obtain the raw intensity signal. The normalized intensity autocorrelation function is computed from the acquired signal and fitted to the semi-infinite domain solution of Correlation Diffusion Equation (CDE) to estimate the blood flow index (*αD_B_*).

The transistor–transistor logic output from the SPAD was acquired using a data acquisition board (NI PCIe-6612, National Instruments, USA) operating in edge counting mode. Photon arrival events were sampled at 20 MHz to obtain the raw temporal intensity fluctuations from which the intensity autocorrelations were computed using an integration time of 3 s. A single detection channel was used throughout all experiments.

The optical probe used for light delivery and collection is shown in Fig. 1. During all measurements, the incident optical power at the tissue surface was maintained at 15 mW over a beam diameter of approximately 3.5 mm, which is below the maximum permissible exposure limit specified by the ANSI Z136.1 laser safety standard. The source and detector fibers were embedded within a custom-fabricated polydimethylsiloxane (PDMS) mold and secured inside a 3D-printed probe housing. Elastic straps enabled stable probe placement and minimized fiber tip movement during data acquisition.

### 2.2 Brief theory of DCS and data acquisition strategy

Photon arrival events detected by the SPAD were recorded continuously for an integration period of 3 s. The resulting photon count fluctuations were used to calculate the normalized intensity autocorrelation function, *g*_2_(*τ*), using a multi-tau autocorrelation algorithm [23]. The normalized electric-field autocorrelation function, *g*_1_(*τ*), was subsequently obtained from *g*_2_(*τ*) using the Siegert relation, *g*_2_(*τ*) = 1+*β*|*g*_1_(*τ*)|^2^, where *β* is the coherence factor determined by the detection optics.

The measured *g*_1_(*τ*) was fitted to the analytical solution of the Correlation Diffusion Equation (CDE) for a semi-infinite homogeneous medium to estimate the Blood Flow Index (BFI), defined as *αD_B_*. The CDE is given by

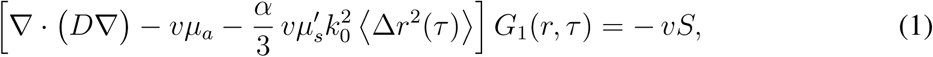

where *D* is the photon diffusion coefficient, *v* is the speed of light in the medium, *µ_a_* and *µ_s_′* are the absorption and reduced scattering coefficients respectively, *k*_0_ = 2*π/λ* is the wave vector where *λ* = 785nm, and ⟨Δ*r*^2^(*τ*)⟩ represents the mean square displacement of the scattering particles. For all analyses, optical properties were fixed at *µ_a_* = 0.01 mm*^−^*^1^ and *µ_s_′* = 1 mm*^−^*^1^ as determined by our in-house developed time domain near infrared spectroscopy system.

The Brownian diffusion model, ⟨Δ*r*^2^(*τ*)⟩ = 6*D_B_τ,* was assumed for the motion of red blood cells, where *D_B_*is the effective diffusion coefficient. The BFI thus obtained was plotted in real time to visualize the live blood flow changes during a certain task protocol.

The photon counting and multi-tau algorithms were implemented in C, while the live acquisition interface and CDE fitting codes were developed in MATLAB R2021b (MathWorks, USA). A 3 s acquisition window was followed by autocorrelation computation and model fitting, resulting in an effective temporal resolution of 8.8 s.

### 2.3 Human study

All experimental procedures involving human participants were approved by the Institutional Ethics Committee (IEC) of the Indian Institute of Technology Bombay under protocol number IITB-IEC/2024/10. The study was conducted in accordance with the principles of the Declaration of Helsinki. Written informed consent was obtained from all participants prior to their enrollment.

Healthy adults aged 18–40 years were recruited for the study. Exclusion criteria included a history of stroke, neurodegenerative disorders, neurological conditions such as epilepsy, recurrent migraine, and any musculoskeletal or neurological impairment affecting the lower limbs that could interfere with the experimental tasks.

#### 2.3.1 Task protocol

The ACA supplies blood to the medial aspect of the frontal and parietal lobes, including the lower-extremity representation of the primary motor cortex. Consequently, lower-limb motor activity is expected to increase metabolic demand within the ACA territory and elicit a corresponding increase in CBF through neurovascular coupling [24, 25]. Therefore, lower-limb motor tasks were selected as physiological perturbations for evaluating the sensitivity of the proposed DCS approach to ACA blood flow changes.

#### 2.3.2 Selection of Optimal Source-Detector Separation

To determine the optimal SD separation for sampling ACA blood flow, measurements were acquired using three SD separations: 15 mm, 17 mm, and 20 mm. For all measurements, the detector position was maintained at the glabella.

Subjects were instructed to perform a standing leg-marching task for 1 min at each SD separation. During the task, participants marched in place while following an metronome set to 70 beats per minute to maintain a certain pace. Consecutive measurements were separated by a 30-min rest period to allow cerebral hemodynamics to return to baseline. The experimental protocol is shown in Fig. 2(a). Five healthy volunteers (3 females and 2 males; age (25 ± 3.7) years) participated in this experiment.

**Fig 2.**
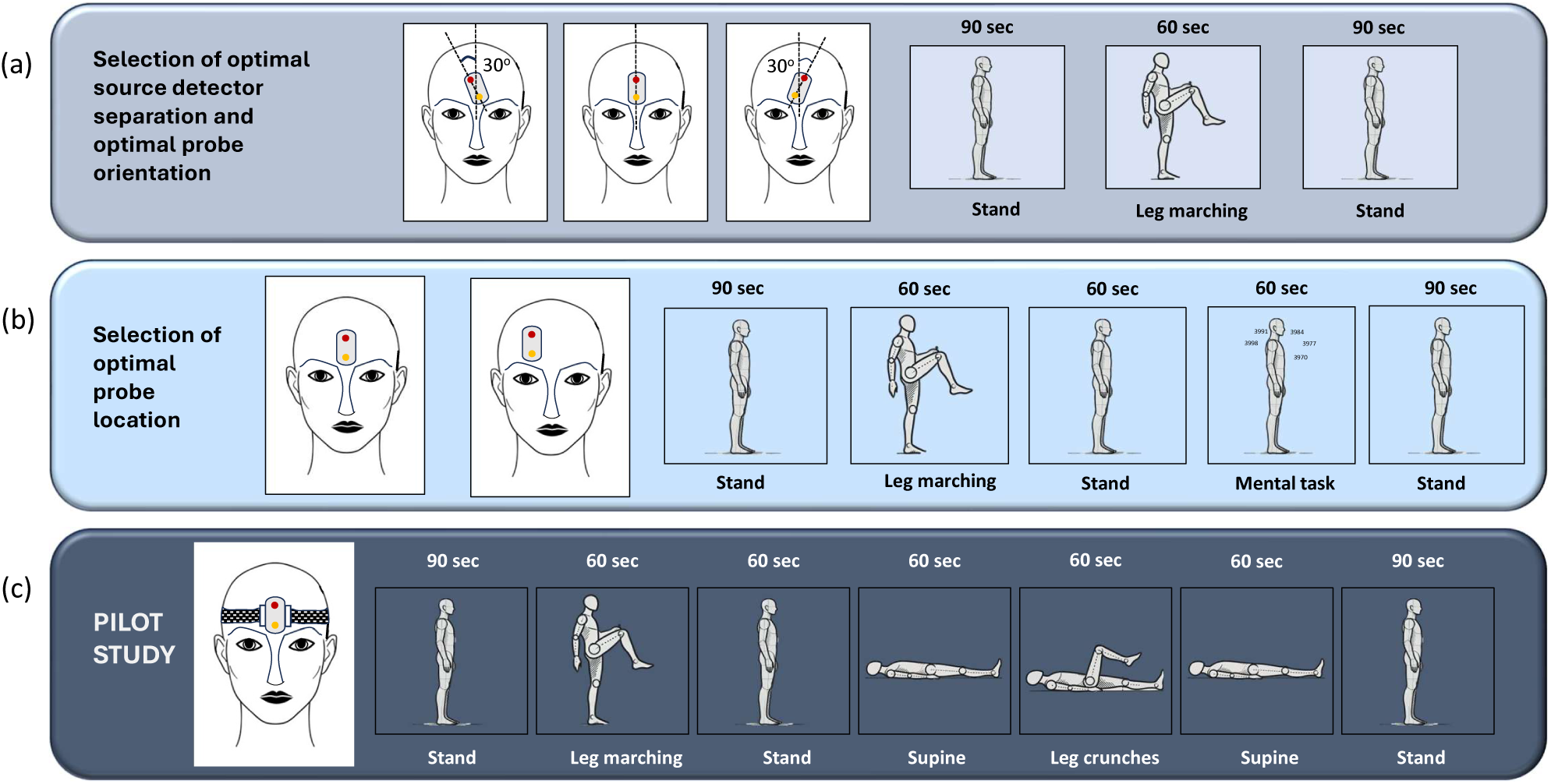
Task protocol for the optimization of probe configuration and placement and the pilot study involving healthy human subjects: Schematic showing the (a) orientation of the probe on glabella about the midline and the corresponding task protocol. (b) Position of the probe on glabella and Fp1 and the corresponding task protocol. (c) Final probe placement with the optimized probe parameters (orientation : vertical (0*^◦^*); placement : glabella) and location and the pilot study protocol.

#### 2.3.3 Selection of Optimal Probe Orientation

Following determination of the optimal SD separation, the effect of probe orientation on ACA sensitivity was investigated. Measurements were acquired at three probe orientations relative to the cranial midline: 30*^◦^* anti-clockwise, vertical 0*^◦^*, and 30*^◦^* clockwise. For each orientation, subjects performed the standing leg-marching task for 1 min. Consecutive measurements were separated by 30-min rest periods. The same subject group from the preceding optimization study participated in this experiment as well.

#### 2.3.4 Selection of Optimal Probe Location

To identify the optimal location for ACA monitoring, DCS measurements were acquired from two frontal locations: the glabella and Fp1. Two perturbation tasks were chosen: a leg-marching task, expected to enhance the blood flow in the paracentral lobule, which is specifically supplied by ACA, and therefore in turn in ACA itself [18]. The second was a mental task, involving serial subtraction by seven from a randomly selected four-digit number, which is known to enhance the blood flow in multiple regions of the cortex involving both frontal and the parietal cortices which are only partly supplied by ACA. [26, 27].

Each participant performed both tasks twice, once with the probe positioned at the glabella and once at Fp1 as shown in Fig. 3(b). Consecutive measurements were separated by a 30-min rest period. Data were acquired from 12 participants. Two datasets were excluded from further analysis. In one case, the participant performed the mental task aloud, introducing speech-related artifacts into the measurements. In the second case, excessive baseline fluctuations prevented reliable assessment of task-induced blood flow changes. Consequently, data from 10 participants (5 females and 5 males; age (27 ± 5.2) years) were included in the final analysis.

**Fig 3.**
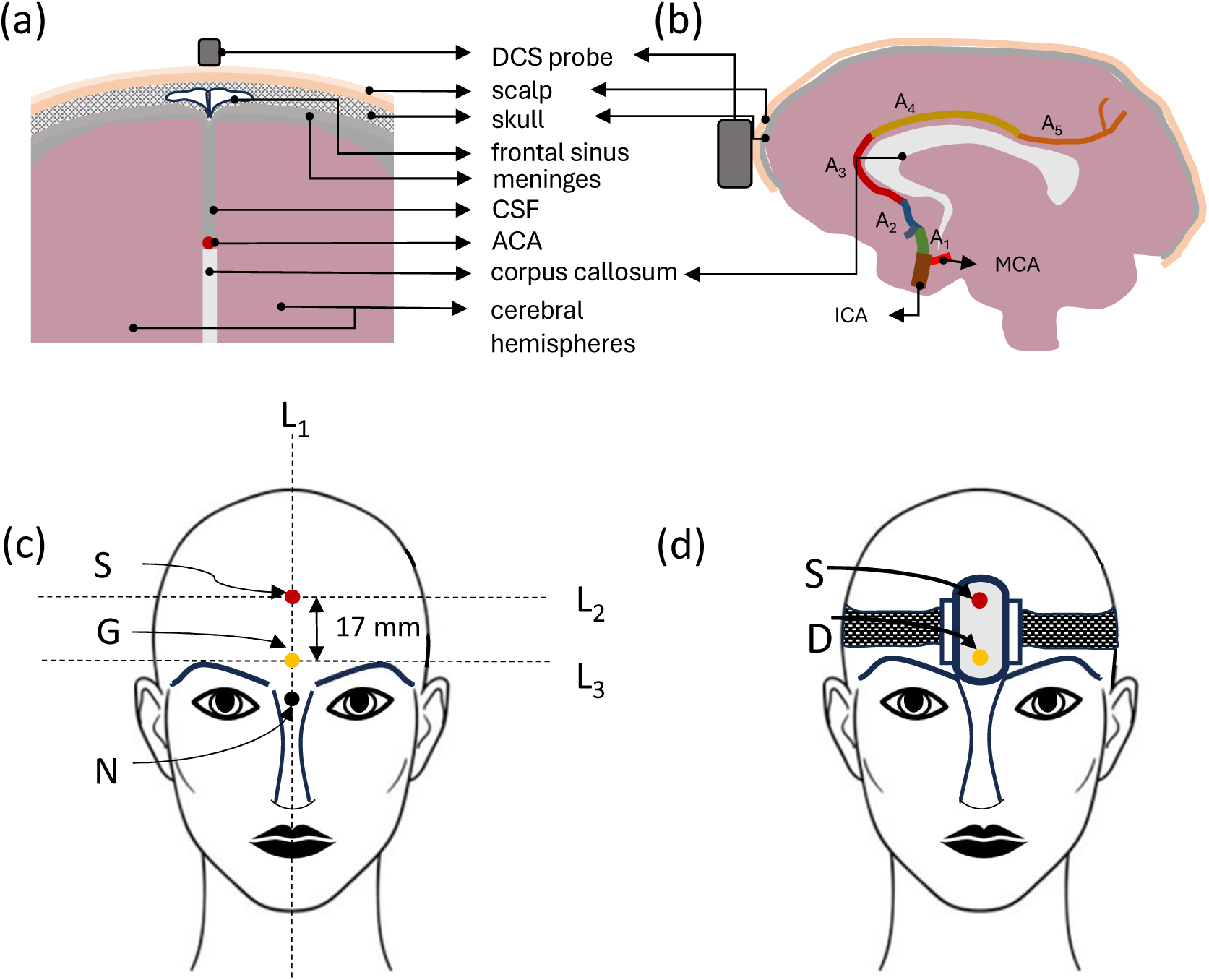
Probe placement for sampling flow from the ACA: (a) Schematic of axial section of brain showing scalp and all layers of the brain up to the ACA. (b) Schematic of sagittal section of the brain showing bifurcation of the Internal Carotid Artery (ICA) into the Middle Cerebral Artery (MCA) and the five segments of ACA - A1, A2, A3, A4 and A5. (c) Schematic (anterior view) of the anatomical landmarks used for DCS probe positioning to target the ACA. N - nasion; G-glabella; L_1_-facial midline; L_2_ and L_3_-lines perpendicular to L_1_ and separated by 17 mm; S and G denote the locations of source and detector fiber-tips respectively, defined by the intersections of L_1_ with L_2_ and L_3_. (d) The final DCS probe placement containing the source (S) and detector (D) positions.

#### 2.3.5 Probe Placement on the Glabella

Based on the results of the SD separation, probe orientation, and location optimization experiments, a standardized probe placement protocol was developed for targeting the blood flow changes in ACA. The final configuration comprised an SD separation of 17 mm, vertical probe orientation, and placement centered on the glabella.

The targeted portion of ACA, comprising of partly the A2 and A3 segments, corresponds to the anterior interhemispheric portion of the artery that courses around the genu of the corpus callosum. From the frontal aspect, this segment represents the first major intracranial artery encountered along the optical path in the longitudinal fissure, thereby maximizing sensitivity to ACA blood flow as shown in Figs. 3(a) and 3(b).

The anatomical landmarks used for probe placement to target the ACA are shown in Fig. 3(c). The nasion N is the midpoint of the frontonasal suture at the junction of the frontal and nasal bones, visible as a midline depression above the nasal bridge and between the eyes. The glabella G is located 7 mm superior to N along the facial midline (L_1_) [28]. An imaginary line, L_3_, passes through G and is perpendicular to L_1_, while L_2_ is parallel to L_3_ and positioned 17 mm above it.

The detector fiber tip (D) is placed at G, corresponding to the intersection of L_1_ and L_3_, as shown in Fig. 3(d), with a placement tolerance of ±2 mm. The source fiber tip (S) is positioned at the intersection of L_1_ and L_2_, resulting in an SD separation of approximately 17 mm.

#### 2.3.6 Phantom Validation

To evaluate the feasibility of detecting blood flow from a vessel located at a depth equivalent to that of ACA from the scalp surface, a multilayer tissue-mimicking phantom was fabricated based on the anatomical thicknesses of the scalp, skull, frontal sinus and ACA.

The phantom consisted of a scalp layer (5 mm, *µ_a_* = 0.019 *mm^−^*^1^, *µ_s_′* = 0.86 *mm^−^*^1^) placed above a skull-mimicking layer (13 mm, *µ_a_* = 0.014 *mm^−^*^1^, *µ_s_′* = 0.66 *mm^−^*^1^) containing a frontal sinus cavity. The depth of the sinus (maximum depth = 13 mm) were obtained from a representative CT slice of an adult human brain as shown in Fig. 4(a). Beneath these layers at a depth of 45 mm below the scalp surface, a tube of diameter 2.5 mm was placed in an intralipid solution which mimicked the optical properties of the Cerebrospinal Fluid (CSF) (*µ_a_* = 0.004 *mm^−^*^1^, *µ_s_′* = 0.003 *mm^−^*^1^). The tube simulated the anterior interhemispheric segment of the ACA [22, 29], through which an aqueous solution of intralipid and India ink to mimic the optical properties of blood (*µ_a_* = 0.54 *mm^−^*^1^, *µ_s_′* = 1.3 *mm^−^*^1^) was circulated using a syringe pump at a flow rate of 5 mL/min. All of the above-mentioned optical properties were obtained from previously reported literature values [30–33].

**Fig 4.**
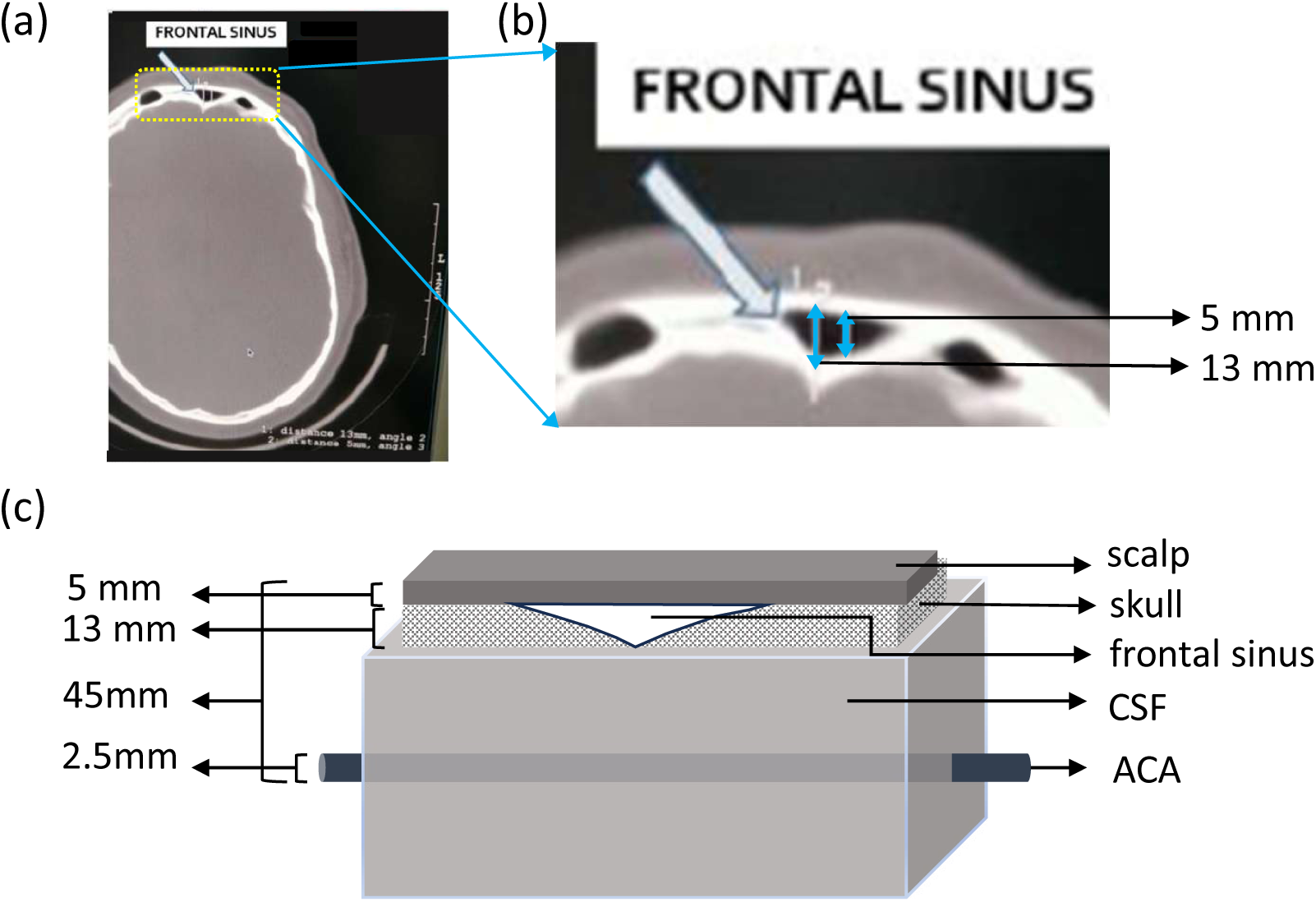
Phantom schematic: (a) A CT scan slice of an adult human brain showing the skull with the dimensions of the frontal sinus. (b) Enlarged view of the frontal sinus and its dimensions. (c) Schematic showing the multilayer phantom where the optical properties and the depth of the layers of the brain up to the ACA have been mimicked.

Using the optimized probe configuration determined from the human studies, the probe was positioned directly above the flow channel with the source and detector aligned parallel to the vessel axis and separated by 17 mm. DCS measurements were recorded continuously throughout the experiment.

To assess the influence of the frontal sinus and the probe position on flow sensitivity, the experiment was repeated once by placing the probe off-center to the vessel axis and then upon replacing the skull layer containing the sinus cavity with a homogeneous solid PDMS block of identical thickness and optical properties. The resulting flow sensitivity was compared between the three cases.

#### 2.3.7 Pilot Study

Following optimization of the probe geometry and validation using the multilayer ACA phantom, the final probe configuration was evaluated in a larger cohort of healthy volunteers. Participants performed two lower-limb motor tasks: (i) standing leg marching and (ii) supine leg crunching interspaced with some postural changes involving supine and standing positions. The standing leg-marching task was included because of its expected activation of the paracentral lobule which is supplied by the A4 segment of ACA. The supine leg-crunch task was incorporated to provide an alternative perturbation for subjects who may have difficulty performing prolonged standing tasks. The complete experimental protocol is shown in Fig. 2(c). A total of 40 healthy volunteers comprising 20 females (age 28.8±7.7 years) and 20 males (age 27.7±7.9 years) participated in the pilot study. An equal number of male and female participants was ensured to achieve a balanced sex representation and avoid potential bias.

### 2.4 Data Analysis

For each subject, the relative Cerebral Blood Flow or rCBF was quantified with respect to the baseline which is the initial standing position as

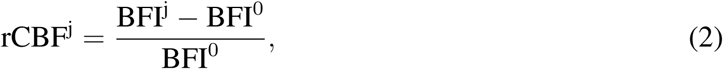

where BFI^j^ is the BFI corresponding to the j*^th^* timestamp and BFI^0^ is the mean BFI of the initial standing position which is the baseline. Similarly the mean rCBF is defined as mean of the rCBF values obtained from Eq. (2) for a given task (leg march or leg crunch or mental task) or position (stand or supine).

### 2.5 Statistical Analysis

To visualize the distribution of rCBF changes across subjects, box-and-whisker plots were generated for the optimization experiments (optimal source-detector separation and probe orientation) and the pilot study. For the optimization experiments, box-and-whisker plots were constructed for the leg-marching and mental tasks during both the task and resting periods. For the pilot study, box-and-whisker plots were generated for the leg-marching task and the standing and supine conditions. In each box, the central line represents the median rCBF value across all subjects. The lower and upper edges of the box correspond to the 25th and 75th percentiles, respectively, defining the interquartile range (IQR). The whiskers extend to the most extreme observations within 1.5 × IQR from the lower and upper quartiles. Data points beyond the whiskers are displayed individually as outliers (red stars).

To determine the significance of the changes observed during tasks the corresponding mean rCBF values were considered which were initially tested for normality using the Kolmogorov–Smirnov test and cross-checked by quantile-quantile plots. Since the data did not satisfy the assumption of normality, paired Wilcoxon signed-rank test, a non-parametric statistical method, was used to assess the significance. For the probe location optimization experiment, comparisons were made between different tasks measured at the same probe location (e.g. leg marching versus mental task at the glabella) and between the same task measured at different probe locations (e.g. leg marching at the glabella versus Fp1). For the pilot study, comparisons were performed between the leg-marching task and the standing condition, and between the supine leg-crunch task and the supine resting condition. All statistical tests were two-sided, and statistical significance was defined as p < 0.01. The analyses were performed using MATLAB R2021b (MathWorks, USA).

The minimum sample size required for detecting task-induced changes in rCBF was determined by an a priori power analysis using G*Power 3.1 software. The analysis was conducted assuming a significance level of *α* = 0.01, a statistical power of 1 − *β* = 0.95, and an expected effect size estimated from probe orientation optimization results for vertical orientation, d=3.9 (average change in rCBF during leg task 73% and a standard deviation of 18%), resulting in a sample size of 5. To ensure adequate representation of both sexes, a minimum sample size of 10 participants was considered necessary. However, a total of 40 participants (20 males and 20 females) were recruited to increase the statistical power of the study findings.

## 3 Results

### 3.1 Optimal Parameter Selection for Pilot Study

#### 3.1.1 Selection of optimal source-detector separation

The effect of SD separation on sensitivity to task-induced CBF changes was evaluated using SD separations of 15 mm, 17 mm, and 20 mm. Representative rCBF obtained by performing the task protocol as shown in Fig. 2(a) from five subjects are shown in Fig. 5(a). While all three separations exhibited an increase in rCBF during the leg-marching task, measurements acquired at 20 mm showed higher variation during both the resting and task periods compared to those obtained at 15 mm and 17 mm.

**Fig 5.**
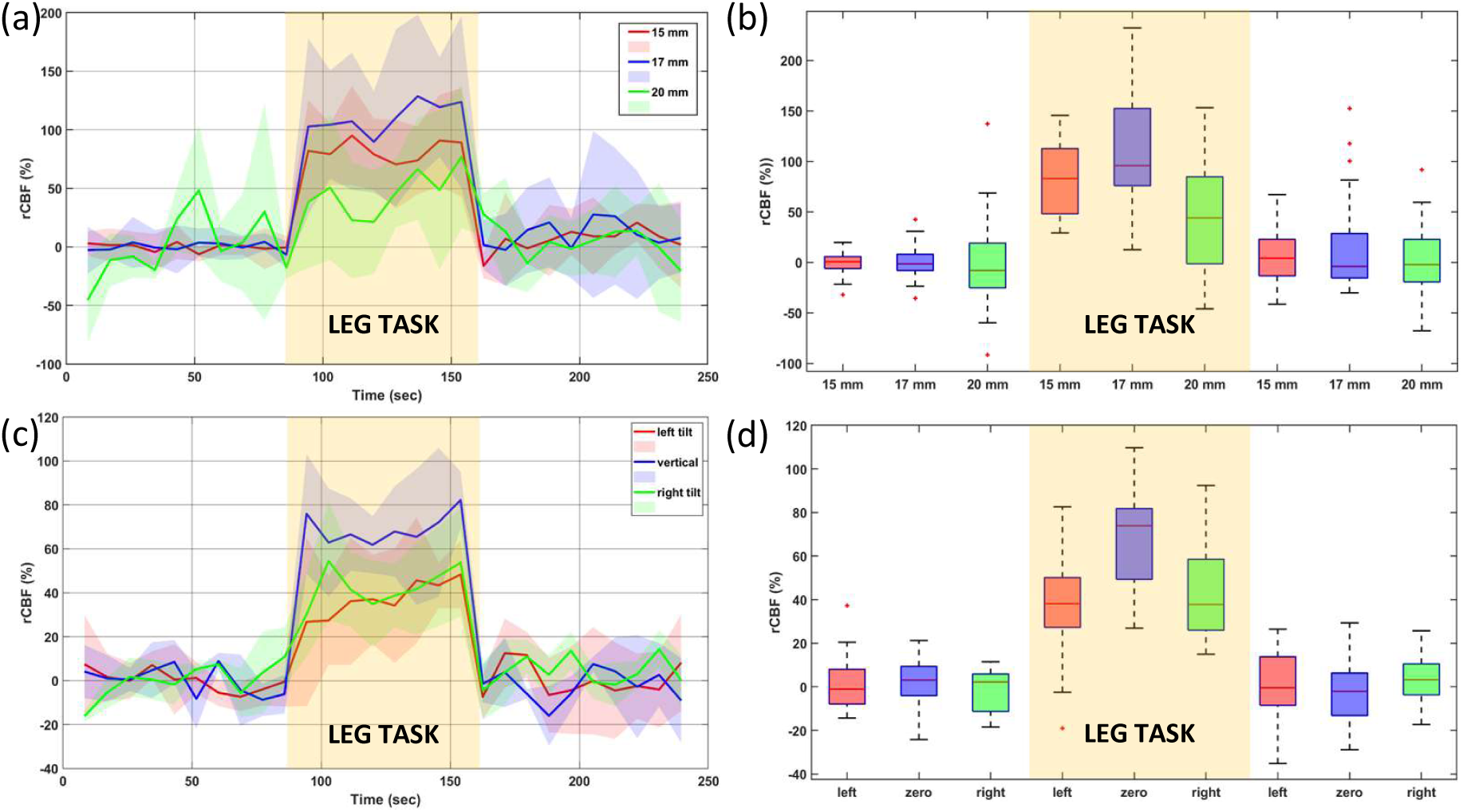
Results for selection of optimal probe configuration for sampling ACA in five subjects: (a) Live rCBF plots for optimal SD selection. (b) Box-whisker plots when the SD separation is 15 mm (red), 17 mm (blue) and 20 mm (green). (c) Live rCBF plots for optimal probe orientation selection. (d) Box-whisker plots when the probe is placed on the glabella in a vertical orientation (blue), tilted anticlockwise by 30*^◦^* (red) and tilted clockwise by 30*^◦^* (green) with respect to the cranial midline.

The distribution of mean rCBF values during the leg-marching task is shown in Fig. 5(b). The median rCBF increase was 83.0% (IQR: 48.1-112.6%) for the 15 mm separation, 95.8% (IQR: 75.9-152.4%) for the 17 mm separation, and 44.0% (IQR: -1.3 to 84.7%) for the 20 mm separation. Among the three configurations, the 17 mm separation produced the largest task-induced increase in rCBF with lower variability than the 20 mm configuration. Therefore, 17 mm was selected as the optimal SD separation for subsequent experiments.

#### 3.1.2 Selection of an optimal probe orientation

Following optimization of the SD separation, the effect of probe orientation was investigated while maintaining the SD separation at 17 mm. Figure 5(c) shows rCBF plots corresponding to the three probe orientations evaluated at 30*^◦^* anti-clockwise tilt, 0*^◦^* vertical orientation, and 30*^◦^* clockwise tilt with respect to the cranial midline. The task protocol for the same is shown in Fig. 2 (a).

The corresponding distributions of mean rCBF values are shown in Fig. 5(d). The median rCBF increase during the leg-marching task was 38.2% (IQR: 27.3-50.1%) for the anti-clockwise orientation, 73.9% (IQR: 49.3-81.8%) for the vertical orientation, and 37.8% (IQR: 26.0-58.5%) for the clockwise orientation. The vertical orientation showed the largest task-induced increase in rCBF and was therefore selected for all subsequent measurements.

#### 3.1.3 Selection of optimal probe location

The final optimization experiment compared measurements acquired from the glabella and Fp1 locations, during the tasks as shown in Fig. 2 (b), using the optimized SD separation (17 mm) and vertical probe orientation.

For measurements acquired at the glabella, both tasks showed increases in rCBF relative to baseline as seen in Figs. 6(a) and 6(b). However, the leg-marching task produced a larger response (51.6 ± 22.5%) than the mental task (17.9 ± 18.9%). In contrast, measurements acquired at Fp1 showed comparable mean increases in rCBF during the leg-marching task (27.2 ± 24.9%) and the mental task (29.9± 25.7%), with a high standard deviation indicating large inter-subject variability as seen in Figs. 6(c) and 6(d).

**Fig 6.**
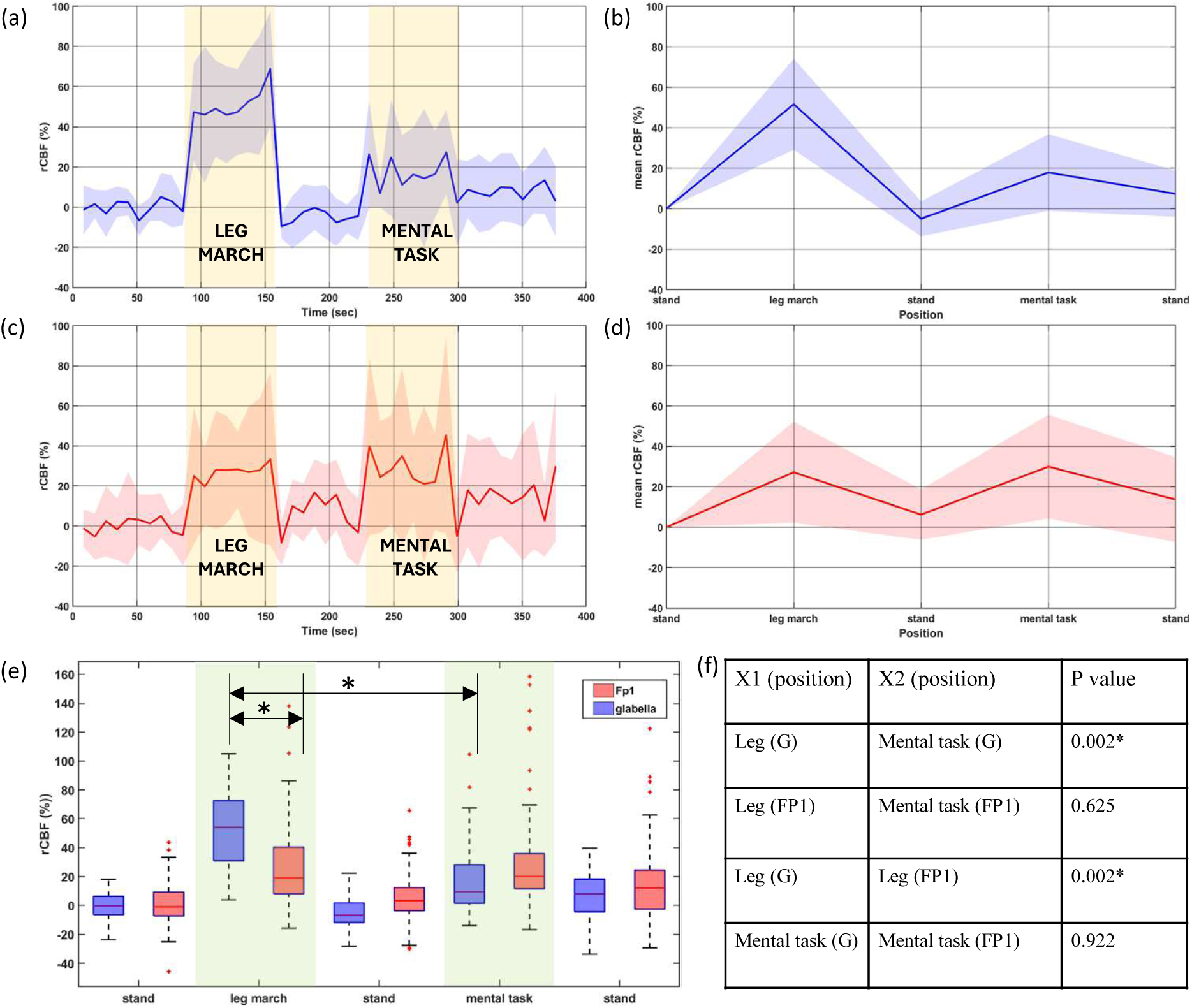
Results for optimal positioning of the probe : (a) Live rCBF plot (bold line - mean, shaded region - standard deviation) across ten subjects when the probe is placed on the glabella. (b) Mean rCBF plot. (c) Live rCBF plot when the probe is placed on Fp1. (d) Mean rCBF plot. (e) Box-whisker plot of the rCBF values from each task and position block from glabella (blue) and Fp1 (red). (f) Table showing the p-values obtained for a pair of task (* indicates p-value < 0.01).

Direct comparison between the two measurement locations revealed that the leg-marching task produced significantly greater rCBF changes at the glabella than at Fp1 (p <0.01) as shown in Figs. 6(e) and 6(f). No significant difference was observed between the two locations during the mental task (p = 0.922). Furthermore, at the glabella, the rCBF increase during leg marching was significantly greater than that observed during the mental task (p < 0.01), whereas no such difference was observed at Fp1 (p = 0.625).

These findings indicate that measurements acquired at the glabella show enhanced sensitivity to hemodynamic changes associated with lower-limb motor activation and are therefore more likely to reflect blood flow changes originating from the ACA territory. Consequently, the glabella was selected as the optimal measurement location.

Based on the above optimization studies, the final probe configuration consisted of an SD separation of 17 mm, a vertical probe orientation, and placement centered on the glabella.

#### 3.1.4 Phantom validation results

The optimized probe configuration was subsequently evaluated using the multilayer ACA-mimicking phantom, as described in Sec. 2.3.6. Discrete parts of the phantom including the scalp, skull with sinus cavity and the ACA-mimicking tube are shown in Fig. 7(a). A top view of the phantom, with all the layers stacked including scalp (5 mm), skull with sinus cavity (15 mm thick), ACA-mimicking tube (located approximately 45 mm below the scalp surface), and an intralipid-water solution medium to mimic the optical properties of CSF, is shown in Fig. 7(b).

**Fig 7.**
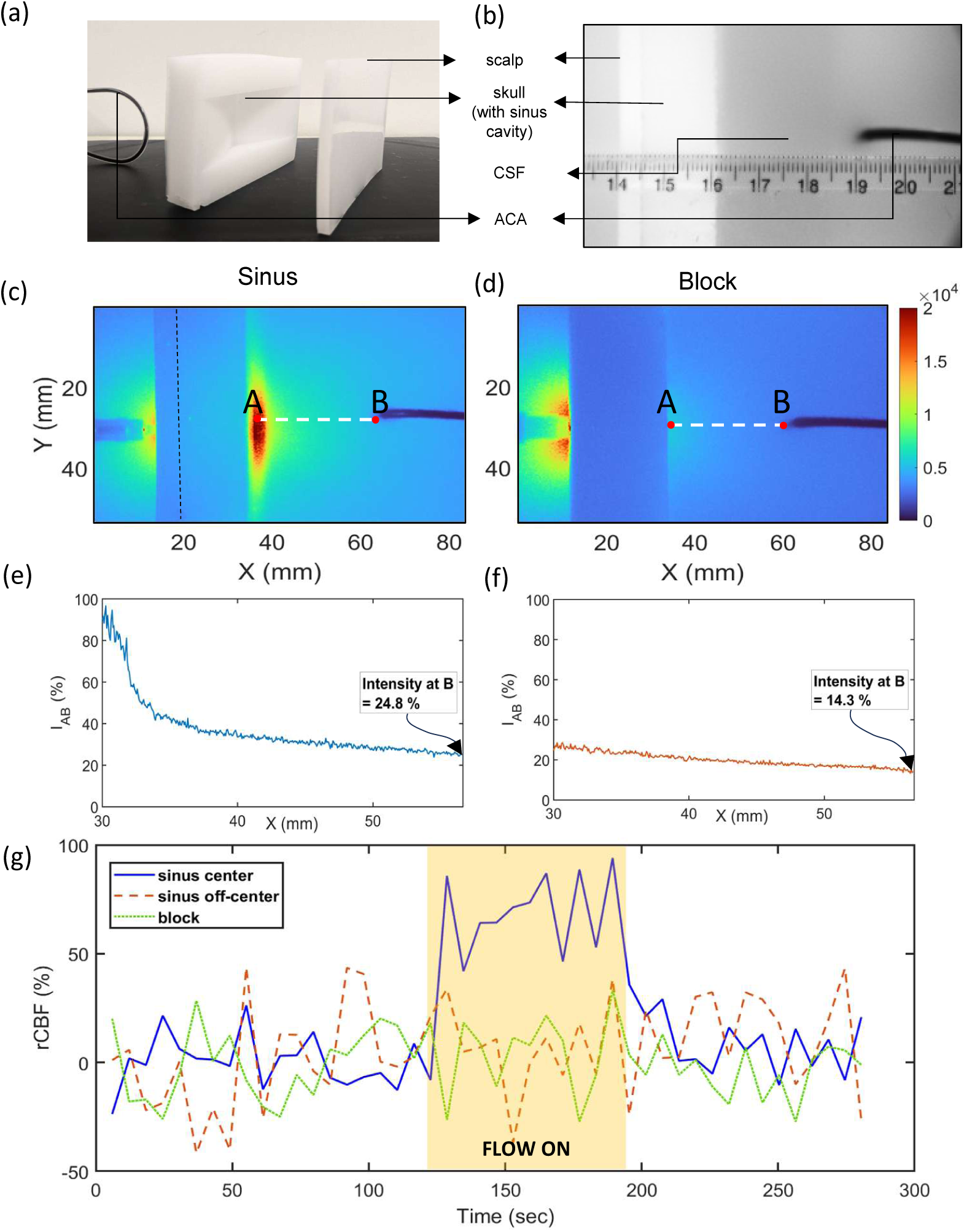
Flow results from ACA mimicking phantom: (a) Image showing the parts of the multilayer phantom - scalp, skull (with the sinus cavity) and the ACA mimicking tube (b) Top view of the multilayer phantom when the layers are stacked in the order shown in (a). (c) Transmitted light profile through the sinus-cavity and (d) the solid-block configurations respectively (e) Transmitted light intensity profile plots measured along the white dotted line AB (I_AB_) up to the tip of the ACA-mimicking tube for the sinus-cavity and (f) the solid-block configurations, respectively. (g) Relative flow measurements for three probe positions: aligned vertically over the sinus cavity and along the tube axis (blue), positioned off-center with respect to the tube axis (red), and placed over the solid block with no sinus cavity (green). The yellow shaded region indicates the interval during which the tube had flow.

To investigate the influence of the frontal sinus on photon propagation the transmitted light distribution was taken for two cases : (i) skull layer containing a sinus cavity and (ii) solid skull without a sinus cavity mimicked by a PDMS block of identical thickness and optical properties, as shown in Figs. 7(c) and 7(d), respectively. The point source was placed at the center of the scalp/block in line with the axis of the tube where the optical power was kept at 15 mW matching the conditions used in the human experiments. The corresponding intensity images were acquired by a scientific CMOS camera at an exposure time of 2 ms. A localized higher intensity region was seen on the incident side in the block case as compared to the sinus case which could be due to the absence of a cavity resulting in higher backscattering. A correspondingly higher intensity region was observed immediately beneath the sinus cavity, suggesting higher photon transmission through the skull with sinus cavity as compared to the solid block case. To quantitatively compare light transmission through these two configurations, intensity along the white-dotted line AB (normalised with respect to the incident intensity) were compared, as shown in Figs. 7(e) and 7(f). The measured intensity at the tip of the tube, as denoted by point B in Figs. 7(c) and 7(d), was found to be higher for the sinus-cavity (24.8%) than for the solid-block configuration (14.3%). These observations indicate that in the presence of a sinus cavity the effective attenuation decreases and the corresponding transmitted light intensity reaching the tube increases as opposed to the solid block case indicating absence of sinus.

We next evaluated the sensitivity of the DCS system to the flow changes within the ACA-mimicking tube where the SD separation was fixed at the 17 mm. Three experimental conditions were investigated: (i) the probe positioned directly above and aligned with the axis of the tube in the multilayer phantom containing the sinus cavity, (ii) the probe positioned off-axis relative to the tube in the same multilayer phantom, and (iii) the probe positioned above the tube in the solid-block configuration without the sinus cavity. During the flow on period the flow rate within the tube was maintained at 5 mL/min. The corresponding DCS measurements are shown in Fig. 7(g), where the baseline is considered as the period before the flow was started.

A clear increase in the measured flow was observed only when the probe was aligned with the tube axis in the presence of the sinus cavity. In contrast, negligible changes were observed for both the off-axis configuration and the solid-block configuration. These findings indicate that the optimized probe geometry provides sensitivity to flow changes in a vessel located approximately 45 mm beneath the scalp surface and that the frontal sinus plays an important role in probing the flow from such deep vessels from the surface.

#### 3.1.5 Pilot study results

The optimized probe configuration was subsequently evaluated in a cohort of 40 healthy volunteers. The rCBF plots are shown in Fig. 8.

**Fig 8.**
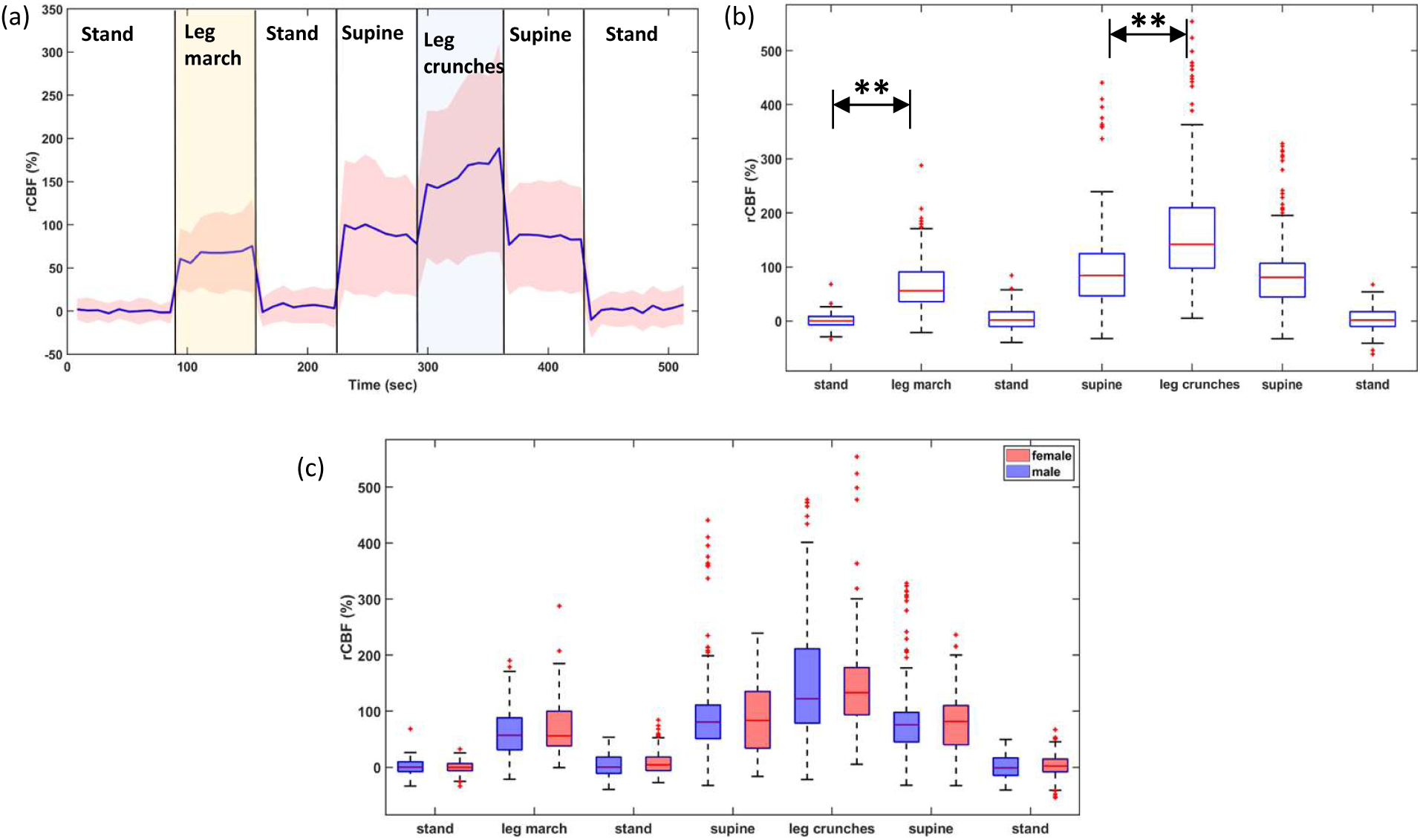
Pilot study results : (a) rCBF plot showing live blood flow changes obtained from the glabella with SD 17 mm and vertical orientation across forty subjects. (b) Box whisker plots showing the distribution of the rCBF values across all subjects for each task/position block. (c) Box-whisker plots showing the rCBF distribution across all male (blue) and female (red) subject groups for each task/position block. (** indicates p-value < 0.001)

The postural changes from standing to supine after the leg marching task resulted in a mean increase in rCBF of 91.6 ± 68.7% and a decrease of 85 ± 59.7% from supine to standing after the leg crunches task. The standing leg-marching task resulted in a mean increase in rCBF of (66.4 ± 38.6%, p<0.001) relative to the standing baseline. Similarly, the supine leg-crunch task produced a mean increase in rCBF of (39.4 ± 32.2%, p<0.001) relative to the supine position.

Figure 8(c) shows the box-and-whisker plots of task-induced rCBF changes for male and female participants. During the standing leg-marching task, the median rCBF increase was 57.1% (IQR: 31.1–88.2%) in males and 56.0% (IQR: 38.2–99.9%) in females. During the supine leg-crunch task, the median rCBF increase was 24.6% (IQR: 5.7–55.7%) in males and 30.4% (IQR: 13.4–51.0%) in females. Overall, the rCBF changes during the postural changes and the standing leg-marching task were comparable between males and females, while females exhibited a slightly higher rCBF response than males during the supine leg-crunch task.

Overall, these results demonstrate that the optimized DCS configuration is capable of detecting task-induced blood flow changes in ACA across a large cohort of healthy subjects.

## 4 Discussion

In the present work, we used DCS for the targeted measurement of blood flow in the ACA, specifically its A3 segment which courses around the genu of the corpus callosum. The A3 segment and its distal branches which form A4 supply the medial frontal cortex and paracentral lobule, regions that contain the lower-limb representation of the motor and sensory homunculus [18]. Consequently, lower-limb motor tasks were employed to evoke localized increases in blood flow within the ACA territory.

A systematic optimization strategy involving SD separation, probe orientation, and probe location was used to maximize sensitivity to blood flow changes in the ACA. The optimization experiments demonstrated that an SD separation of 17 mm, a vertical probe orientation, and probe placement at the glabella produced the largest task-induced rCBF responses. Furthermore, the glabellar location showed significantly greater sensitivity to ACA-related lower-limb motor activation than to the mental task, which primarily increases blood flow in the prefrontal cortex. This observation suggests enhanced sensitivity from glabella to blood flow changes originating from the ACA. In addition, the rCBF response during the mental task was higher at Fp1 than at the glabella, although the variance was also considerably larger. This increased variability may be attributed to greater contamination from extracerebral tissues due to the thicker skull in the prefrontal region. This finding is consistent with the fact that the dorsolateral prefrontal cortex (Brodmann areas 9 and 46), which is primarily engaged during numerical processing tasks, lies closer to the Fp1 location [26, 27]. Although the ACA also supplies portions of these regions along with MCA, its contribution is comparatively smaller, resulting in only a small increase in rCBF at the glabella during the mental task [18]. Sensitivity to ACA-specific blood flow changes could be further improved through the use of filtering techniques designed to reduce extracerebral contamination in deep-tissue flow measurements [34].

A crucial factor contributing to the high depth sensitivity achieved from the glabella is the absence of cortical tissues in the longitudinal fissure. As shown in Fig. 4(a), the only optically attenuating medium between the skull and the ACA is the CSF, which has very low optical properties (*µ_s_′* = 0.003 mm*^−^*^1^ and *µ_a_* = 0.004 mm*^−^*^1^). Combined with the frontal sinus cavity, this results in higher light intensity reaching the ACA. Consequently, photons scattered from the ACA were able to reach the detector with sufficient sensitivity to detect flow changes, as shown in Fig. 7. We note that the flow rate in our setup was limited to 5 mL/min due to the technical constraints of the syringe pump, which is substantially lower than the physiological blood flow rate in the ACA of an adult human brain (82 ± 18 mL/min) [35]. Despite this limitation, a clear increase in the flow was detected in the sinus case as compared to the off-axis probe placement and the solid-block configuration, which mimics the absence of the sinus cavity. These findings provide a possible anatomical explanation for the successful detection of ACA blood flow changes from the glabellar region despite the deep location of the vessel. In contrast, regions just off the midline (away from the longitudinal fissure) contain cortical tissues with high optical attenuation (*µ_s_′* = 1.2 mm*^−^*^1^ and *µ_a_* = 0.016 mm*^−^*^1^) which would restrict sensitivity to primarily the superficial cortical layers.

In the pilot study, a significant increase in rCBF was observed during both lower-limb motor tasks. However, the magnitude of the increase varied considerably across subjects, as illustrated by the box-whisker plot in Fig. 8(b). Differences in local metabolic demand and neurovascular coupling may lead to inter-subject variability in ACA blood flow responses, despite the task pace being standardized at 70 bpm using a metronome. Furthermore, the differences in rCBF between the standing and supine conditions exhibited substantial variability, likely due to the relatively short resting periods between tasks. Although longer recovery intervals would have allowed for more complete physiological stabilization and potentially reduced variability, they would also have considerably increased the overall duration of the protocol, making it less practical for participant compliance. Additionally, no statistically significant sex-related differences were observed in the task-induced rCBF responses during either the standing leg-marching task (p = 0.14) or the supine leg-crunch task (p = 0.35).

An important implication of this work is the potential application of DCS for continuous bed-side monitoring of ACA perfusion. Current non-invasive approaches for assessing cerebral blood flow in large intracranial vessels rely primarily on TCD. The major drawbacks of TCD include variability in acoustic windows across patients, the requirement for favorable insonation geometry, and dependence on a skilled operator [5]. In practice, ACA measurements are most commonly performed through the transtemporal acoustic window, where the dominant signal originates from the middle cerebral artery (MCA), while reliable assessment of distal ACA segments remains challenging [4, 6]. In contrast, DCS can be continuously deployed at the bedside using a lightweight optical probe and does not require acoustic access through the temporal bone. The ability to monitor ACA-related blood flow changes from the glabella may therefore provide a complementary tool for cerebral perfusion monitoring in acute neurological conditions, including ischemic stroke. Apart from bedside monitoring, targeted measurement of ACA blood flow may have important applications in post-stroke care. Continuous or repeated measurements of task-evoked rCBF could potentially provide longitudinal information regarding perfusion status within the ACA territory. Such measurements may be useful for tracking the temporal evolution of cerebral perfusion following stroke and for assessing responses to therapeutic interventions. However, validation in patient populations will be required before such applications can be realized.

Despite these promising results, several limitations should be considered. Anatomical variability of the ACA may influence measurement sensitivity. Variations in vessel trajectory, depth, curvature, and interhemispheric positioning can alter the relationship between the optical sampling volume and the targeted ACA segment [29, 36]. In some individuals, a larger or smaller SD separation, or a slight deviation from the vertical probe orientation, may provide improved sensitivity. Similarly, inter-subject differences in scalp thickness, skull morphology, frontal sinus anatomy, and cortical structure may influence photon propagation and the optimal measurement geometry. Consequently, the parameters identified in this study should be considered optimal for the general population rather than subject-specific. Future studies combining Magnetic Resonance Angiography (MRA) with DCS measurements could provide individualized probe placement and improve sensitivity to ACA blood flow changes. A second limitation of the current system is its temporal resolution. Although the system was capable of resolving task-induced blood flow changes, the effective temporal resolution of 8.8 s limits the ability to monitor rapid hemodynamic fluctuations. Further optimization of the acquisition pipeline, autocorrelation computation, and fitting algorithms may improve temporal resolution.

To the best of our knowledge, this is the first study demonstrating the use of DCS for targeted measurement of blood flow within a specific intracranial artery through optimization of probe geometry and placement. The primary objective of this work was to establish the feasibility of non-invasive ACA monitoring using a systematic combination of SD separation optimization, probe orientation optimization, and anatomical positioning at the glabella. Rather than quantifying absolute ACA blood flow or comparing DCS measurements against established vascular imaging modalities, our focus was on demonstrating that reproducible ACA-sensitive blood flow changes could be detected using a practical bedside-compatible optical system. The optimization experiments, phantom validation studies, and large-cohort pilot measurements collectively provide evidence supporting the feasibility of this approach. Future studies should focus on validating the technique against established vascular imaging methods, improving subject-specific probe placement strategies, and evaluating its utility for continuous cerebral perfusion monitoring in patients with acute cerebrovascular disorders.

## 5 Conclusion

This study showed the feasibility of targeted, non-invasive measurement of blood flow in the ACA using DCS. Through systematic optimization of SD separation, probe orientation, and probe location, a glabellar measurement configuration was identified that showed consistent sensitivity to ACA-related blood flow changes during lower-limb motor activation tasks. The optimized configuration was further supported by anatomically relevant phantom experiments and validated in a cohort of 40 healthy volunteers, where significant increases in rCBF were observed during two independent leg-based tasks. The proposed approach extends the capability of DCS beyond conventional cortical perfusion measurements towards targeted monitoring of specific intracranial vascular territories and may provide a practical framework for continuous bedside assessment of ACA perfusion. Such targeted measurements could facilitate longitudinal monitoring of cerebrovascular status and provide physiologically relevant data for neuromonitoring, and aid in future clinical decisions.

## Disclosures

H Varma, U S Srinivasan, S Das, K Sharma, S Sarkar and K Murali are the inventors on relevant patents. The authors declare that there are no financial interests, commercial affiliations, or other potential conflicts of interest that could have influenced the objectivity of this research or the writing of this paper.

## Code, Data, and Materials Availability

The code and data utilized in this study are available from the authors upon reasonable request.

## Acknowledgments

This research was funded by Wadhwani Research Centre of Bioengineering, Indian Institute of Technology Bombay (DO/2025-WRCB002-108), Technology Innovation Hub, Indian Institute of Technology Bombay (RD/0125-TIH0018-005), Core Research Grant, Science and Engineering Research Board (RD/0122-SERB000-049), HDFC ERGO (DO/2024-HDRD002-007).

## Biographies

**Susweta Das** received her B.Sc. and M.Sc. degrees in Physics from Calcutta University, Kolkata, and Delhi University, New Delhi, India, in 2017 and 2019 respectively. She is currently pursuing her Ph.D. degree in Biomedical Engineering at Indian Institute of Technology, Bombay. Her research focuses on developing laser speckle-based cerebral blood flow monitoring systems and supporting algorithms aiming towards clinical translation, particularly for bedside monitoring in stroke-affected patients.

**Kavita Sharma** received her M.Sc. degree in Physics from Maharshi Dayanand University, Rohtak, India, in 2017, and her Ph.D. degree in Physics from the Central University of Rajasthan, Ajmer, India, in 2024. She served as a Mentor/Co-Supervisor under the IEEE Photonics Society Internship Program from November 2023 to January 2024. She also worked as a Project Scientist at SeNSE, Indian Institute of Technology Delhi, India, from February 2024 to August 2024. She is currently a Postdoctoral Researcher in the Department of Biosciences and Bioengineering at the Indian Institute of Technology Bombay, India. Her current research focuses on the development of Laser Speckle Correlation Microscopy systems in reflection mode, Raman spectroscopy system development and platform stabilization studies, and cerebral blood flow monitoring in stroke patients, including system performance enhancement and clinical translational studies.

**Soumyajit Sarkar** received B-Tech degree in electrical engineering from West Bengal University of Technology, Kolkata in 2015. He completed his Ph.D. in Biomedical engineering from Indian Institute of Technology-Bombay. His research interest includes the development of models and associated systems to measure time varying cerebral blood flow using laser speckle-based method associated with both clinical and preclinical studies.

**Kimberly Gonsalves** received her B.Tech. degree in Bioengineering from MIT World Peace University, Pune, in 2026. She is currently pursuing research as an intern at the Indian Institute of Technology Bombay, within the Theoretical and Experimental Bioimaging Lab under the guidance of Prof. Hari M. Varma. Her research interests focus on the design and development of optical biomedical imaging systems and the implementation of hardware-accelerated processing architectures for high-speed physiological data acquisition.

**U. S. Srinivasan** received the M.Ch. degree in Neurosurgery from Madurai Medical College, Madurai, India, in 1993. He has over 32 years of experience in neurosurgery and has held several academic and clinical appointments, including establishing the Department of Neurosurgery at PSG Medical College, Coimbatore, serving as Chief Neurosurgeon at MIOT Hospitals, Chennai, and as Consultant Neurosurgeon at Nizwa Hospital, Oman. He is currently Senior Consultant Neurosurgeon at Sri Balaji Hospitals, Chennai, India. He has authored scientific publications and books and collaborates with the Indian Institute of Technology Bombay and the Indian Institute of Technology Madras on the development and clinical evaluation of neurosurgical technologies.

**Hari M. Varma** received the M.Sc. (Engg.) and Ph.D. degrees from the Indian Institute of Science, Bangalore, India, in 2004 and 2010, respectively. He was a postdoctoral fellow in the Department of Mathematics and Systems Analysis at Aalto University, Helsinki, Finland, during 2010 and 2011, and subsequently served as a postdoctoral fellow in the Medical Optics Group at the Institute of Photonic Sciences, Barcelona, Spain, from 2012 to 2015. He is currently an Associate Professor of Biomedical Optics in the Department of Biosciences and Bioengineering at the Indian Institute of Technology Bombay, India. His primary research focuses on developing systems and methods for biomedical optics, particularly speckle-based imaging systems for imaging cerebral blood flow in both animal models and clinical settings.

## List of Figures

1. DCS Schematic
2. All experiment protocol and probe placement
3. Probe placement for ACA
4. Phantom schematic
5. Optimal SD results
6. Optimal location results
7. Phantom results
8. Pilot results

## Notes

### Competing Interest Statement

The authors have declared no competing interest.

## References

[1] S. M. Greenberg, W. C. Ziai, C. Cordonnier, et al., “2022 guideline for the management of patients with spontaneous intracerebral hemorrhage: a guideline from the american heart association/american stroke association,” Stroke 53(7), e282–e361 (2022).

[2] R. C. Nogueira, M. Aries, J. S. Minhas, et al., “Review of studies on dynamic cerebral autoregulation in the acute phase of stroke and the relationship with clinical outcome,” Journal of Cerebral Blood Flow & Metabolism 42(3), 430–453 (2022).

[3] J. Vymazal, A. M. Rulseh, J. Keller, et al., “Comparison of ct and mr imaging in ischemic stroke,” Insights into imaging 3(6), 619–627 (2012).

[4] R. Baumgartner, “Transcranial insonation,” Frontiers of neurology and neuroscience 21, 105 (2006).

[5] M. Marinoni, A. Ginanneschi, P. Forleo, et al., “Technical limits in transcranial doppler recording: Inadquate acoustic windows,” Ultrasound in medicine & biology 23(8), 1275– 1277 (1997).

[6] L. Lennihan, G. Petty, M. Fink, et al., “Transcranial doppler detection of anterior cerebral artery vasospasm.,” Journal of Neurology, Neurosurgery & Psychiatry 56(8), 906–909 (1993).

[7] M. Sloan, E. Haley Jr, N. Kassell, et al., “Sensitivity and specificity of transcranial doppler ultrasonography in the diagnosis of vasospasm following subarachnoid hemorrhage,” Neurology 39(11), 1514–1514 (1989).

[8] T. Durduran and A. G. Yodh, “Diffuse correlation spectroscopy for non-invasive, micro-vascular cerebral blood flow measurement,” Neuroimage 85, 51–63 (2014).

[9] D. A. Boas, L. Campbell, and A. G. Yodh, “Scattering and imaging with diffusing temporal field correlations,” Physical review letters 75(9), 1855 (1995).

[10] C. Udina, S. Avtzi, M. Mota-Foix, et al., “Dual-task related frontal cerebral blood flow changes in older adults with mild cognitive impairment: a functional diffuse correlation spectroscopy study,” Frontiers in Aging Neuroscience 14, 958656 (2022).

[11] T. Durduran, G. Yu, M. G. Burnett, et al., “Diffuse optical measurement of blood flow, blood oxygenation, and metabolism in a human brain during sensorimotor cortex activation,” Optics letters 29(15), 1766–1768 (2004).

[12] Q. Wang, Y. Hua, C. Li, et al., “Real-time diffuse correlation spectroscopy with on-chip correlators for measuring human cerebral blood flow and brain function,” Journal of Innovative Optical Health Sciences (2026).

[13] B. Kim, S. Zilpelwar, E. J. Sie, et al., “Measuring human cerebral blood flow and brain function with fiber-based speckle contrast optical spectroscopy system,” Communications Biology 6(1), 844 (2023).

[14] R. C. Mesquita, S. S. Schenkel, D. L. Minkoff, et al., “Influence of probe pressure on the diffuse correlation spectroscopy blood flow signal: extra-cerebral contributions,” Biomedical optics express 4(7), 978–994 (2013).

[15] C. G. Favilla, R. C. Mesquita, M. Mullen, et al., “Optical bedside monitoring of cerebral blood flow in acute ischemic stroke patients during head-of-bed manipulation,” Stroke 45(5), 1269–1274 (2014).

[16] B. L. Edlow, M. N. Kim, T. Durduran, et al., “The effects of healthy aging on cerebral hemodynamic responses to posture change,” Physiological measurement 31(4), 477–495 (2010).

[17] S. Das, S. Sarkar, U. S. Srinivasan, et al., “Direct non-invasive measurement of blood flow changes in the anterior cerebral artery using diffuse correlation spectroscopy,” in Clinical and Translational Neurophotonics 2025, 13302, 53–56, SPIE (2025).

[18] H. Blumenfeld, “Neuroanatomy through clinical cases, with sylvius 4,” Sunderland, MA: Sinauer Associates (2010).

[19] A. Chandra, W. A. Li, C. R. Stone, et al., “The cerebral circulation and cerebrovascular disease i: Anatomy,” Brain circulation 3(2), 45–56 (2017).

[20] E. Kumral, G. Bayulkem, D. Evyapan, et al., “Spectrum of anterior cerebral artery territory infarction: clinical and mri findings,” European Journal of Neurology 9(6), 615–624 (2002).

[21] S. Y. Kang and J. S. Kim, “Anterior cerebral artery infarction: stroke mechanism and clinical-imaging study in 100 patients,” Neurology 70(24 part 2), 2386–2393 (2008).

[22] F. B. Gomes, M. Dujovny, F. Umansky, et al., “Microanatomy of the anterior cerebral artery,” Surgical neurology 26(2), 129–141 (1986).

[23] D. Magatti and F. Ferri, “Fast multi-tau real-time software correlator for dynamic light scattering,” Applied optics 40(24), 4011–4021 (2001).

[24] R. Aaslid, “Visually evoked dynamic blood flow response of the human cerebral circulation.,” Stroke 18(4), 771–775 (1987).

[25] A. A. Phillips, F. H. Chan, M. M. Z. Zheng, et al., “Neurovascular coupling in humans: physiology, methodological advances and clinical implications,” Journal of Cerebral Blood Flow & Metabolism 36(4), 647–664 (2016).

[26] L. Rueckert, N. Lange, A. Partiot, et al., “Visualizing cortical activation during mental calculation with functional mri,” Neuroimage 3(2), 97–103 (1996).

[27] P. Burbaud, O. Camus, D. Guehl, et al., “A functional magnetic resonance imaging study of mental subtraction in human subjects,” Neuroscience letters 273(3), 195–199 (1999).

[28] V. Vigo, K. Cornejo, L. Nunez, et al., “Immersive surgical anatomy of the craniometric points,” Cureus 12(6) (2020).

[29] S. Pai, R. Kulkarni, and R. Varma, “Microsurgical anatomy of the anterior cerebral artery-anterior communicating artery complex: an indian study,” Neurology Asia 10, 21–28 (2005).

[30] H. Zhao and E. M. Buckley, “Influence of oversimplifying the head anatomy on cerebral blood flow measurements with diffuse correlation spectroscopy,” Neurophotonics 10(1), 015010–015010 (2023).

[31] Q. Wang, M. Pan, Z. Zang, et al., “Quantification of blood flow index in diffuse correlation spectroscopy using a robust deep learning method,” Journal of Biomedical Optics 29(1), 015004–015004 (2024).

[32] J. Selb, D. A. Boas, S.-T. Chan, et al., “Sensitivity of near-infrared spectroscopy and diffuse correlation spectroscopy to brain hemodynamics: simulations and experimental findings during hypercapnia,” Neurophotonics 1(1), 015005–015005 (2014).

[33] N. Bosschaart, G. J. Edelman, M. C. Aalders, et al., “A literature review and novel theoretical approach on the optical properties of whole blood,” Lasers in medical science 29(2), 453–479 (2014).

[34] R. Paul, K. Murali, and H. M. Varma, “High-density diffuse correlation tomography with enhanced depth localization and minimal surface artefacts,” Biomedical optics express 13(11) (2022).

[35] L. Zarrinkoob, K. Ambarki, A. Wåhlin, et al., “Blood flow distribution in cerebral arteries,” Journal of Cerebral Blood Flow & Metabolism 35(4), 648–654 (2015).

[36] S. Kedia, S. Daisy, K. K. Mukherjee, et al., “Microsurgical anatomy of the anterior cerebral artery in indian cadavers,” Neurology India 61(2), 117–121 (2013).

